# A combination of systemic mannitol administration and mannitol-modified polyester nanoparticles facilitate gene delivery to the brain through caveolae-mediated endocytosis

**DOI:** 10.1101/2024.08.29.610249

**Authors:** G Betsy Reshma, Chirag Miglani, Arundhati Karmakar, Sarika Gupta, Asish Pal, Munia Ganguli

## Abstract

Overcoming the blood-brain barrier (BBB) remains a significant challenge for nucleic acid delivery to the brain. We have explored a combination of mannitol-modified poly (β-amino ester) (PBAE) nanoparticles and systemic mannitol injection for crossing the BBB. We incorporated mannitol in the PBAE polymer for caveolae targeting and also selected monomers that may help avoid delivery to the liver. We also induced caveolae at the BBB through systemic mannitol injection in order to create an opportunity for the caveolae-targeting nanoparticles (M30 D90)containing plasmid DNA to cross the BBB. When a clinically relevant dose was administered intravenously in this caveolae induction model, M30 D90 demonstrated significant transgene expression of a reporter plasmid in the brain, with selective uptake by neuronal cells and minimal liver accumulation. We also demonstrate that both caveolae modulation using systemic mannitol administration and caveolae targeting using designed nanoparticles are necessary for efficient delivery to the brain. This delivery platform offers a simple, scalable, and controlled delivery solution and holds promise for treating central nervous system diseases with functional targets.

## Introduction

More than 1000 central nervous system (CNS) diseases affect over 1 billion people globally.^[1]^ Nucleic acid drugs with rich target selection present promising therapeutic alternatives for brain disorders that were previously deemed untreatable.^[2]^ Unfortunately, multiple biological barriers prevent efficient nucleic acid delivery into the brain. The blood-brain barrier (BBB), considered the most important one, is a cellular dynamic barrier that protects the CNS from blood components. Barrier properties of the BBB arise from tight junctions between endothelial cells, preventing paracellular movement of molecules, and the low transcytosis rate, preventing intracellular movement of molecules and imposing selective permeability.^[3]^

Several strategies have been developed over the years based on a molecular understanding of the BBB to facilitate the delivery of small and large molecules, viewing the BBB as an interface for delivery.^[4,5]^ Much research focuses on active targeting approaches, with one or more targeting ligands attached to the delivery system to facilitate receptor-mediated transcytosis.^[6,7]^ However, this increases the size and complexity of the delivery system. Moreover, the receptor expression is dynamic with age and disease; limiting the potential of these strategies.^[8–10]^ Another approach is to modulate the BBB permeability with transient tight junction opening strategies. Antibodies and peptides against tight junction proteins^[11,12]^, intracarotid administration of hyperosmotic mannitol^[13]^, and focused ultrasound^[14]^ have been utilized to deliver small molecules. Enhancing transcytosis of the delivery system and modulating the permeability of BBB has worked best for macromolecular cargoes. Studies have shown that pairing delivery systems with focused ultrasound enables efficient transgene expression and genome editing.^[15–17]^

Recent literature suggests that the caveolae-mediated transcytosis pathway across BBB represents a relatively unexplored delivery method for the brain.^[18–20]^ This nonspecific pathway has gained significant attention lately due to observations indicating a transition from clathrin-mediated to caveolae-mediated transcytosis in aging mice.^[10]^ Bacteria^[21]^ and viruses^[22]^ have also been reported to exploit this pathway to cross the BBB. Given that caveolar transport is kept very low at the BBB^[23]^, studies demonstrate that this pathway is agonist inducible with chemical agents such as methamphetamine or physical stimuli like focused ultrasound.^[24,25]^ These studies imply that heightened caveolar transport for a short duration can open up significant opportunities for delivery. It is important to note that the overall permeability increases with caveolae modulators, which might not specifically increase the permeability of the drug or the delivery agent.

To enable nucleic acid delivery, the delivery system must efficiently overcome the poor pharmacokinetics of the nucleic acid.^[26]^ Poly (β-amino ester) (PBAE) based polymeric systems have shown great potential for the delivery of nucleic acid cargo, with their chemical versatility optimized for transfection efficiency and organ targeting.^[27–29]^ While there are no reports of these systems crossing the BBB via systemic routes without additional stimuli like ultrasound^[17]^, there is evidence of safe and efficient delivery when administered locally.^[30–32]^ These nanoparticle-based delivery systems typically use multiple routes to enter the cell in vitro. One way to enhance their selectivity for caveolae-mediated endocytosis is by employing osmotic stimuli.^[33]^ In our previous work, we developed backbone-modified sugar alcohol doped PBAE for caveolae selective entry.^[34]^ We found that mannitol-modified polymeric nanoparticles enhanced transfection efficiency in neuronal cells and enabled selective caveolae-mediated endocytosis. We hypothesized that in an in vivo scenario, such caveolae-selective nanoparticles in the presence of a caveolae-induction model would help nucleic acid delivery across the BBB. Furthermore, we hypothesized that using appropriate monomers for designing the PBAE backbone might avoid delivery to the liver and maximize the chance of reaching the brain.^[35]^

Our manuscript demonstrates that mannitol-modified nanoparticles (M30 D90) carrying plasmid DNA (pDNA) effectively transfect neuronal cells and enhance transport across brain endothelial cells via the caveolae-mediated transcytosis pathway. We explored the use of systemic hyperosmolar mannitol to induce caveolae at the BBB. Although a direct link between hyperosmolar mannitol administration and increased caveolar transport has not been conclusively proven, speculations suggest this relationship.^[36,37]^ Our findings show caveolae puncta in the brain after mannitol injection without compromising the integrity of tight junctions, indicating successful caveolae induction. We show that caveolae abundance is leveraged by M30 D90 enabling efficient transfection in the brain in vivo. Notably, transfection in the liver was low with M30 D90, and uptake was primarily observed in neurons, highlighting the potential for treating central nervous system disorders. In summary, we developed and implemented a simple, efficient, and safe polymer-based delivery system for gene delivery to the brain.

## Results

Based on literature evidence we chose to target caveolae-mediated transcytosis for delivery to the brain. Weinduce caveolae with hyperosmolar mannitol injected systemically. To selectively target the caveolae-rich BBB we built the PBAE system with mannitol modification (Figure 1).

**Figure 1:**
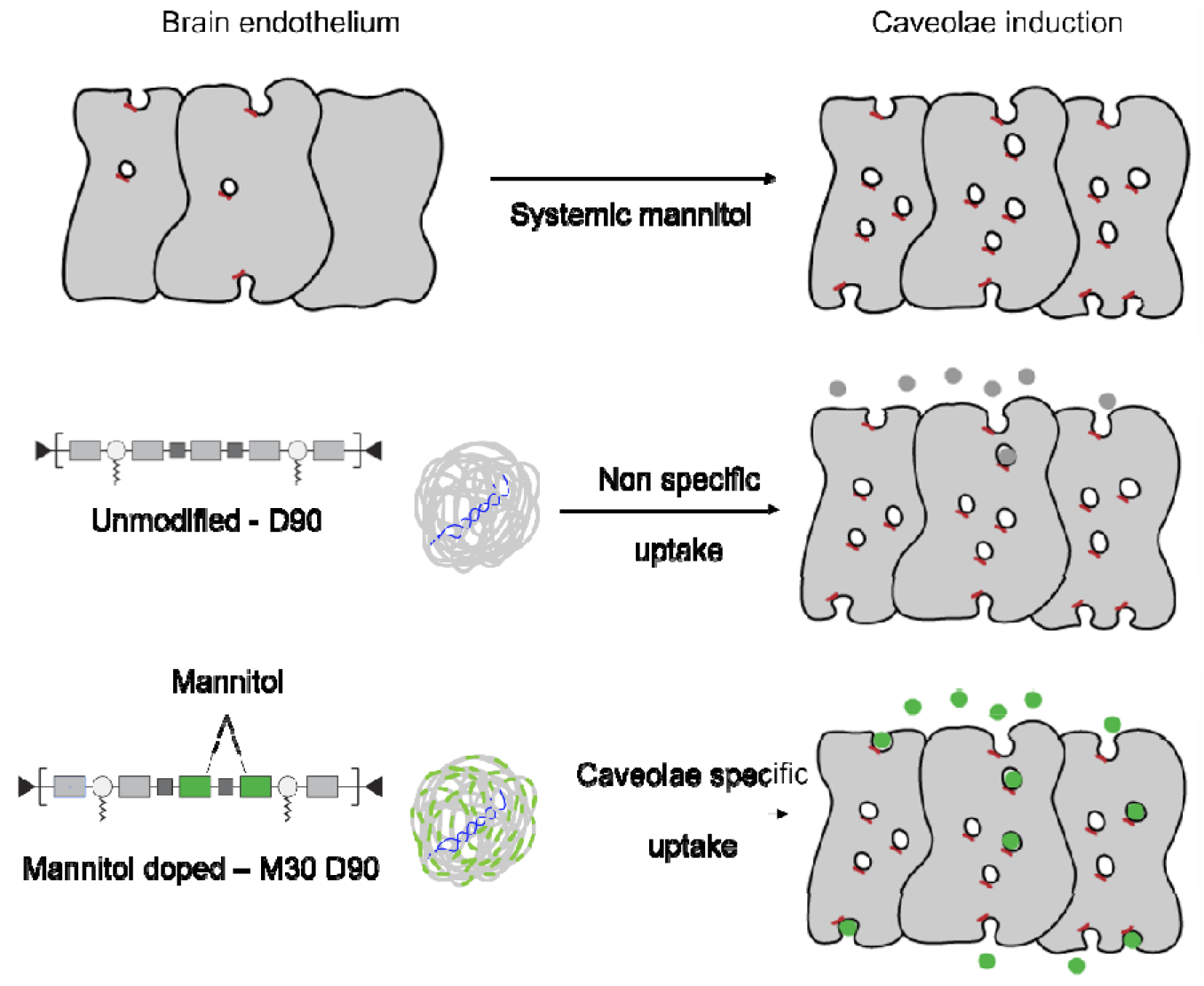
Schematic of the hypothesis suggesting that a combination of caveolae modulation using systemic mannitol along with designing a caveolae selective delivery agent would help in overcoming the blood-brain barrier.

### Synthesis and characterization of mannitol-modified nanoparticles

The PBAE polymer is a combination of diacrylate and amine monomers. We chose Bisphenol A ethoxylate diacrylate (D), reported to escape the liver and have better in vivo transfection of pDNA (plasmid DNA), over Bisphenol A glycerolate diacrylate (DD) which we have used earlier as the diacrylate monomer.^[35]^We used 4-(2-aminoethyl) morpholine (S90) and 1-dodecylamine (Sc12) as the two amine monomers, for the synthesis of the D90 polymer; making it a unique combination not reported so far. For the synthesis of M30 D90, mannitol diacrylate was used to replace D acrylate by 30%, keeping the proportion of the other monomers the same. The exact molar ratios are listed in Table S1. The final polymers of D90 and M30 D90 were obtained by end-capping with diethylenetriamine (E63). The polymer composition and synthesis scheme are depicted in Figure 2a. Proton NMR confirmed 26% doping of mannitol in the backbone of the M30 D90 polymer (Figure S1 and Figure S2). Gel permeation chromatography (GPC) estimated the molecular weight of D90 to be 5082 Da with a PDI of 1.30 and M30 D90 was 5901 Da with a PDI of 1.14 (Table S2). We further measured the nucleic acid binding and osmolarity of the synthesized polymers. M30 D90 had less nucleic acid binding as compared to D90 (Figure 2b) and the osmolarity of M30 D90 was about 130mOsm higher than that of D90 (Figure 2c). M30 D90 also showed a concentration-dependent increase in osmolarity while unmodified D90 did not. Analysis using gel electrophoresis revealed complete complexation of pDNA with the polymers at ratios of 1:5 (pDNA to polymer) for D90 and 1:10 for M30 D90 (Figure S3a). This trend is consistent with the nucleic acid binding data, indicating that D90 binds more efficiently, thus completely encapsulating pDNA with less polymer than M30 D90. Release studies with heparin (anionic challenge) treated nanoparticles (formed at a 1:60 (w/w) of pDNA to polymer) demonstrated complete release even at low heparin amount (1:0.25 w/w polymer to heparin) in the case of M30 D90. Conversely, while release from D90 nanoparticles initiates at lower amounts of heparin, complete DNA release cannot be achieved even with high heparin ratios (1:32) (Figure S3b). Polymeric nanoparticles were formed via self-assembly in aqueous buffer (25mM sodium acetate, pH 5.2) by pipette mixing pDNA and the polymer at a 1:60 (w/w) ratio (pDNA: polymer). D90 and M30 D90 polymers formed nanoparticles with pDNA with a hydrodynamic diameter of 60-70nm (Figure 2d) and polydispersity index less than 0.3 (Figure 2e). Further characterization using Transmission Electron Microscopy (TEM) demonstrated that the morphology was spherical for M30 with pDNA in a ring structure with a hollow center (Figure 2f). The surface charges of D90 and M30 D90 were about +40mV (Figure 2g). Mannitol modification in D90 did not alter the size and charge of the assembled nanoparticles.

**Figure 2:**
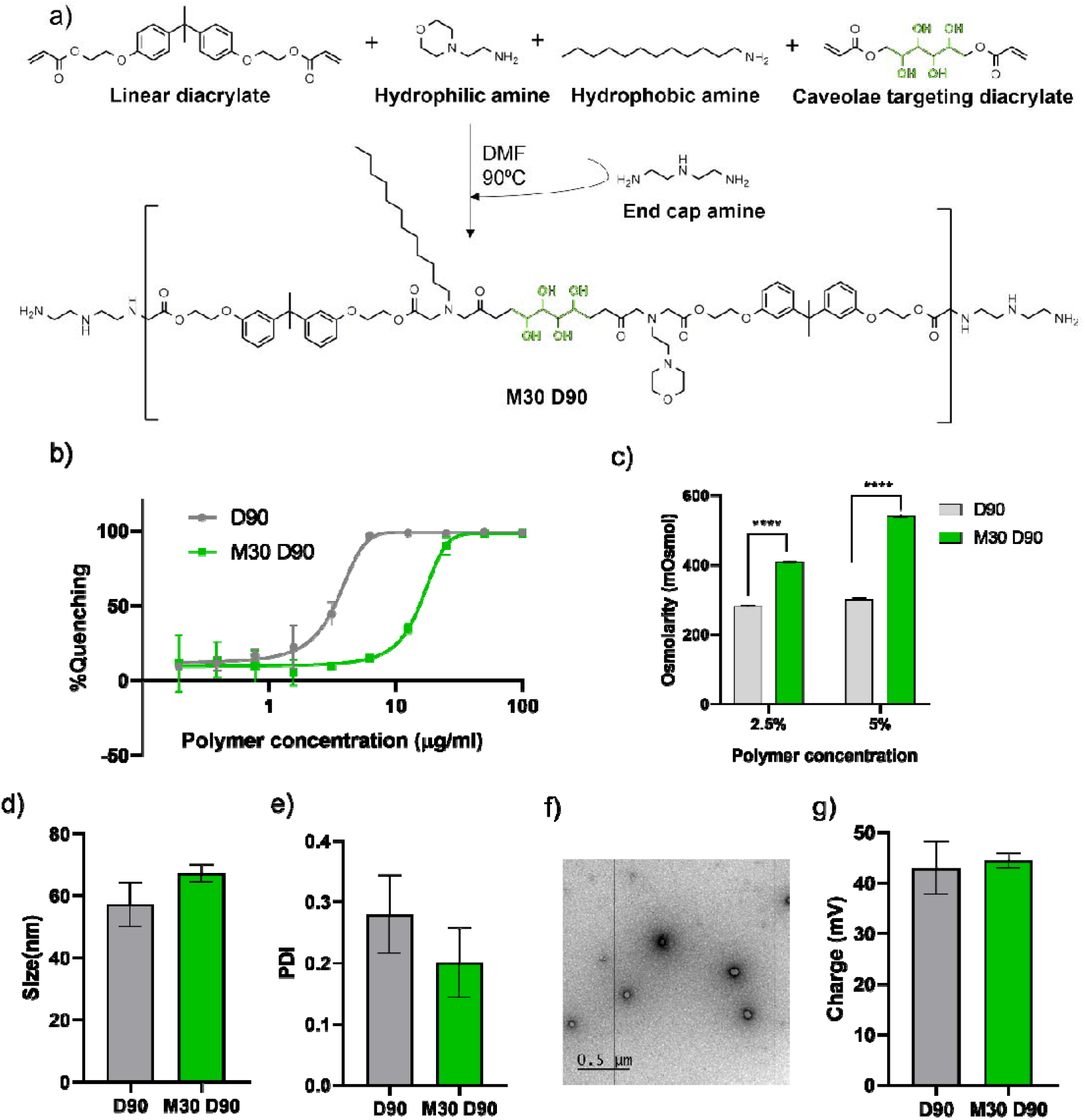
Effects of mannitol modification in D90-Sc12-E63 backbone. a) Scheme of synthesis of mannitol-modified PBAE. Polymer properties b) nucleic acid binding studied with RiboGreen assay and c) Osmolality of polymers. Data is shown as mean ± SD, n=2. Significance was calculated by Sidak’s multiple comparison****p<0.0001. Polymeric nanoparticles were formed with a 1:60 (w/w) ratio of pDNA to polymer in an aqueous buffer. Several nanoparticle properties were measured d) Hydrodynamic size and e) Polydispersity Index estimated by Dynamic Light Scattering. f) Transmission Electron Micrograph of M30 D90 (Scale: 0.5μm) and g) Zeta potential of polymeric nanoparticles. Data represented as mean ± SD, n=3.

### Cellular viability, transfection, and transport of mannitol-modified nanoparticles

The mechanism of uptake of polymeric nanoparticles (where the pDNA is labeled with ATTO488) was checked brain microvascular endothelial cell line (bEND.3)as this endothelial cell type would be the first entry point to the brain in an in vivo scenario. Pretreatment with different endocytotic inhibitors showed different effects was a significant decrease in uptake, more than 30% with nystatin (inhibitor of caveolae-mediated endocytosis); however, there was no notable change in uptake with chlorpromazine (inhibitor of clathrin-mediated endocytosis) or cytochalasin-D (inhibitor of macropinocytosis) for M30 D90 (Figure 3a). This indicates a significant amount of cellular entry of nanoparticles is through the caveolar route although there could be other pathways that we have not explored here. On the other hand, the uptake of D90 nanoparticles did not drop significantly (and in some cases showed an increase) in the presence of endocytosis inhibitors used in this study, suggesting nonspecific or unexplored modes of uptake (Figure 3a). To further validate the involvement of the caveolae-mediated endocytosis pathway, we examined the expression of Caveolin-1 (cav-1), a primary marker for caveolae. We observed overexpression of cav-1 at both transcriptional and translational levels with M30 D90 nanoparticles (Figure 3b and Figure 3c) and not in the case of D90.To check if selectivity to caveolae-mediated endocytosis led to better penetration, we measured the transport of nanoparticles across the endothelial barrier using an in vitro BBB model (Figure 3d). We found approximately a four-fold increase in the transport of M30 D90 across the endothelial cells compared to D90 (Figure 3e). Our findings collectively demonstrate evidence of caveolar involvement in the uptake and improved transport of the M30 D90 nanoparticles(Figure 3f).

**Figure 3:**
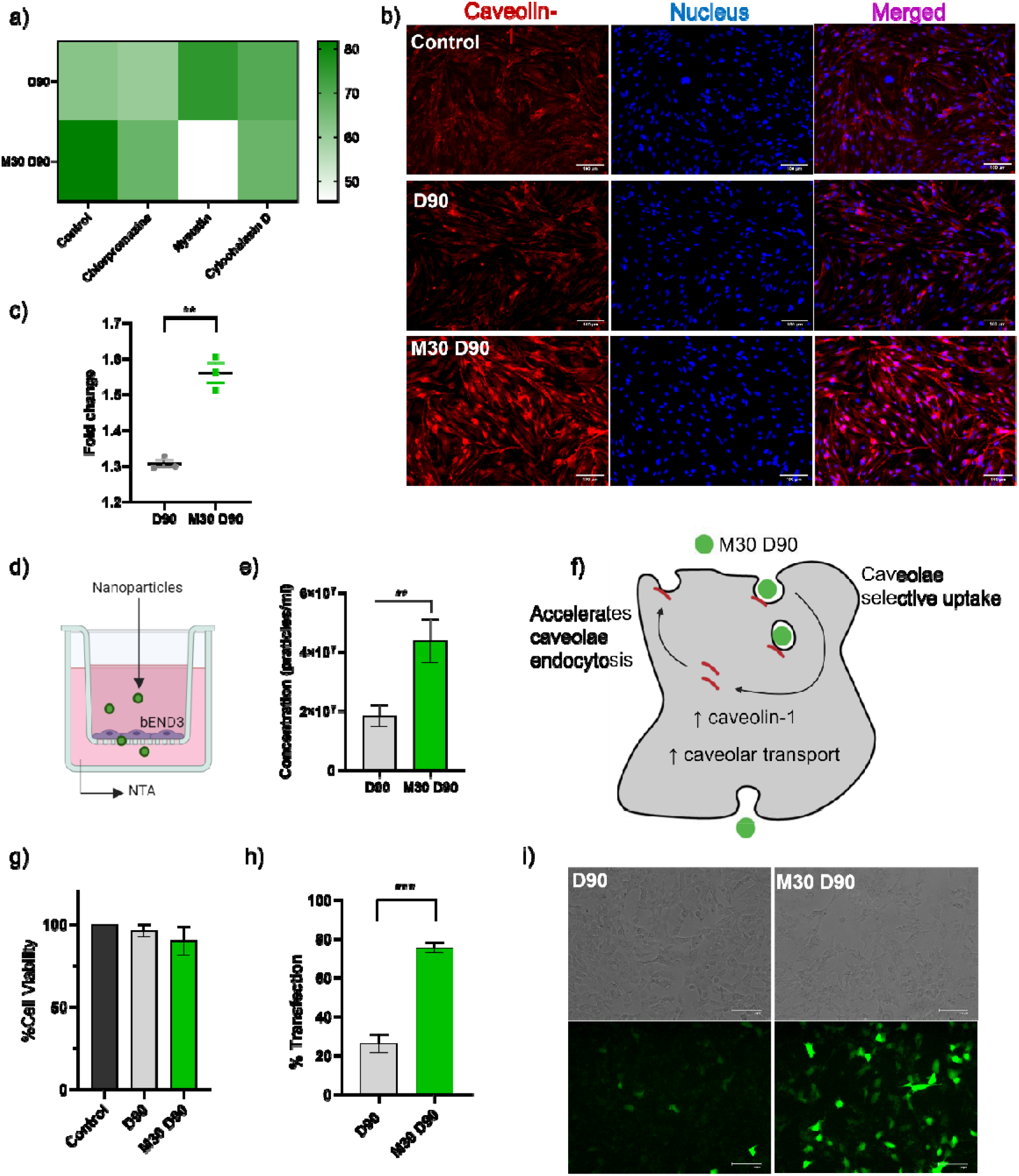
Effects of mannitol modification uptake mechanism, transport, and transfection efficiency. a) Mechanism of endocytosis of nanoparticles (labeled with ATTO488) with pharmacological inhibitors was assessed in brain microvascular endothelial cells (bEND.3). Flow Cytometry analysis revealed a drop in uptake of M30 D90 with nystatin (caveolae inhibitor). b)Immunocytochemistry in brain microvascular endothelial cells (bEND.3) indicating overexpression of cav-1 protein in M30 D90 following 30 minutes incubation with nanoparticles (scale bar: 100μm). c) qRT of cav-1 in bEND.3 after30 minutes of D90 and M30 D90 treatment. Significance was calculated by unpaired t-test correction with Welch’s correction. d) in vitro transwell model of BBB. e) Nanoparticle Tracking Analyzer (NTA) assessment of nanoparticle concentration in media collected from receptor chamber following 5 hours incubation of nanoparticles (labeled with ATTO488) added in the apical chamber. f) M30 D90 is selective for caveolae-mediated uptake, increase cav-1 and increase transport across endothelial cells. Cell viability and transfection efficiency in neuronal cell line (SHSY5Y) with nanoparticles formed with reporter plasmid of green fluorescence protein (pEGFP-C1). g) Percentage cell viability estimated by MTT assay h) Flow cytometry analysis depicting the percentage of transfected cells with M30 D90 compared to D90.i) Fluorescence Microscopy images of transfection (scale bar: 100μm). Data are presented as mean ± SD, n=3, **p < 0.01, ***p < 0.001.

Cellular viability and transfection were assessed in the neuronal cell line SHSY5Y using reporter plasmid pEGFP-C1. M30 D90 and D90 exhibited excellent viability comparable to control, as determined by MTT assay (Figure 3g). Flow cytometry analysis of a low-dose transfection in SHSY5Y cells (300ng per 24 wells) revealed 80% fluorescence-positive cells M30 D90, whereas D90 showed only about 25% fluorescence-positive cells (Figure 3h and 3i). The uptake of M30 D90 was predominantly through caveolae even in the SHSY5Y cell line (Figure S4).

### Effect of systemic mannitol administration on caveolae formation in brain endothelial cells

While M30 D90 increases cav-1 levels in cultured brain endothelial cells, it is unlikely to be adequate for enabling passage across the BBB in vivo. In a healthy brain, non-specific caveolae-mediated endocytosis is kept low.^[38]^ For nanoparticles to utilize this pathway, caveolar transport needs to be induced. Transient upregulation of cav-1 has been reported to increase the transcytosis rate at the BBB^[39]^. Previous studies have demonstrated that systemic administration of hyperosmolar mannitol can increase fluid flux and the formation of pinocytic vesicles in the brain.^[36,40]^ There have been speculations that these pinocytic vesicles are caveolae.^[37]^ Earlier reports have noted increased transgene expression with rAAV-mediated delivery followed by systemic mannitol injection.^[41,42]^ Based on this evidence, we administered hyperosmolar mannitol intraperitoneally to induce caveolae formation at the BBB and observed caveolae puncta marked by cav-1 (Figures 4a and 4b). Systemic administration of mannitol did not compromise the tight junction integrity of the BBB as depicted by unchanged patterns of occludin-1 evidenced through immunostaining (Figure 4c). This finding suggests that mannitol administration induces caveolae and does not compromise the tight junctions of the endothelium. Therefore, we opted to utilize systemic mannitol injection as an agonist for caveolae induction.

**Figure 4:**
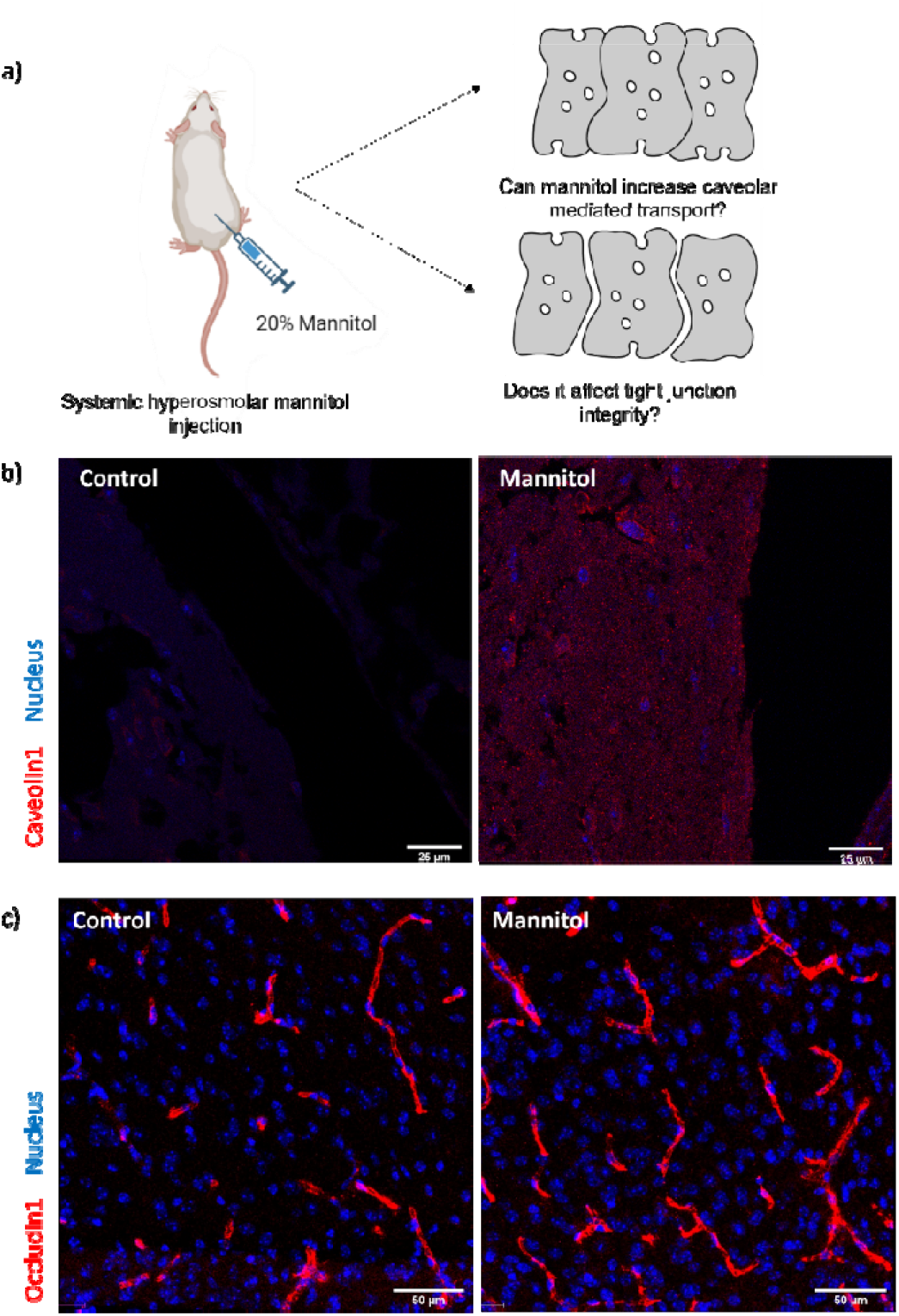
Caveolae can be induced with systemic hyperosmolar mannitol. a)Mannitol-mediated caveolae induction. b) Expression of cav-1 in the brain assessed by immunofluorescence (red) with cell nuclei counterstained with DAPI (blue) (scale bar:25μm) c) Representative images of the tight junction integrity with immunostaining of occludin-1 (red) and nuclei (blue) (scale bar:50μm).

### Effect of mannitol-modified polymeric nanoparticles on in vivo transfection of the brain

Before proceeding in vivo, polyethylene glycol (PEG) was incorporated into the nanoparticle system to enhance stability and reduce interaction with blood components during systemic injection. Alkyl side chains in the polymer backbone allow for the non-covalent incorporation of DMG PEG 2000.^[43]^ The percentage of PEG lipid added was optimized and the volume ratio of buffer and ethanol was adjusted to prepare nanoparticles that are less than 100nm in size as described in the Materials and Methods section (Figure 5a). D90 and M30 D90 with PEG formed uniform nanoparticles with PDI less than 0.3 (Figure S5a) and sizes less than 100nm (Figure 5b). TEM analysis reveals the spherical morphology of assembled M30 D90 nanoparticles with DNA localizing in a peripheral ring structure similar to nanoparticles formulated without PEG (Figure 5c). The size of the PEG-incorporated nanoparticles was similar to that of nanoparticles without PEG. However, there was a significant decrease of over 20mV in the surface charge of the PEGylated nanoparticles, consistent with previous reports of similar systems (Figure 5d).^[28]^ PEG modification did not affect the transfection efficiencies of the polymeric nanoparticles in vitro (Figure S5b).M30 D90 with PEG was stable for up to 48 hours at room temperature as observed with unaltered size and PDI of nanoparticles measured in regular intervals (Figure S5c and S5d).

**Figure 5:**
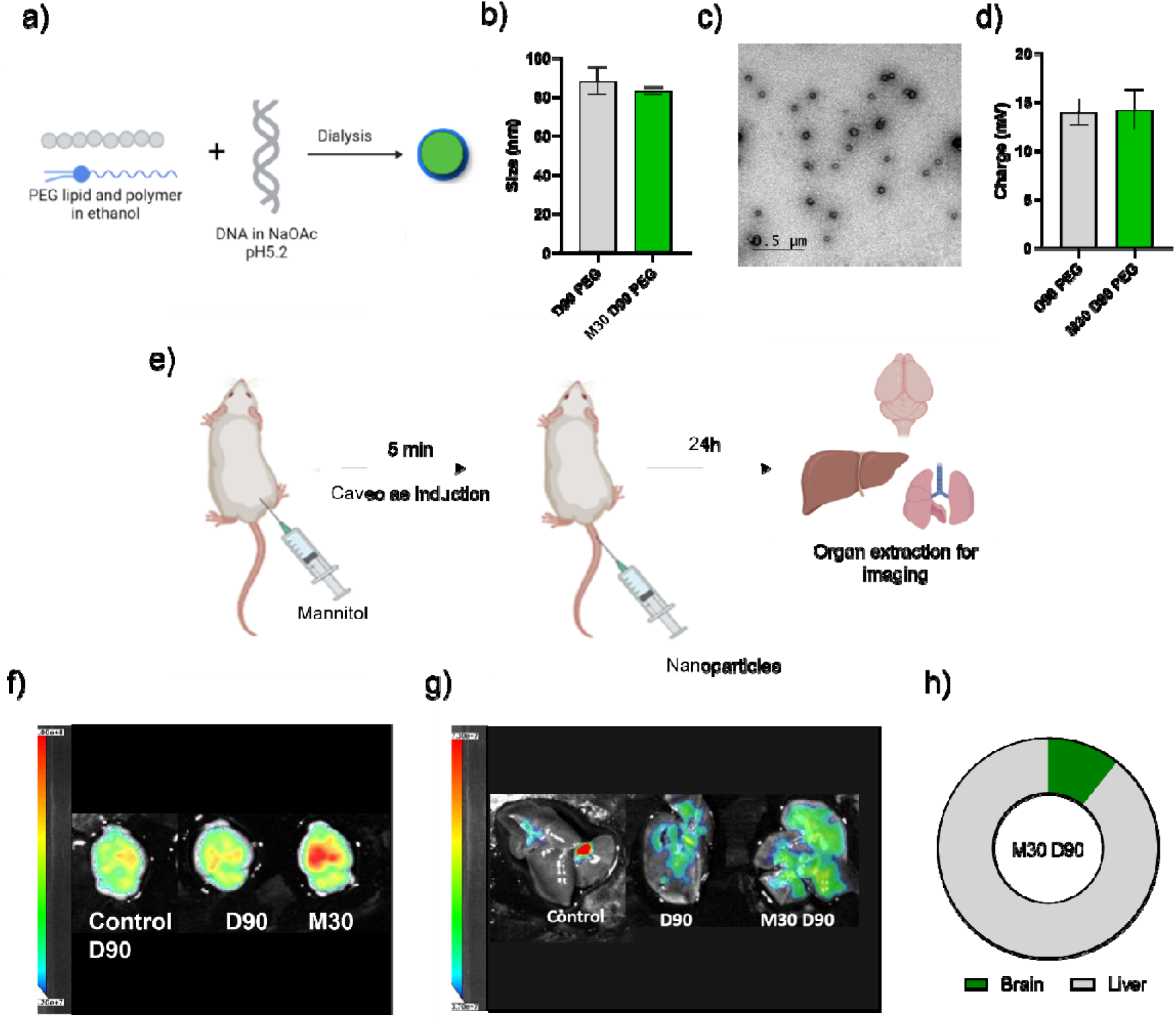
PEG-incorporated M30 D90 transfects the brain following caveolae induction. a) Scheme of PEG lipid incorporation in polymeric nanoparticles. b) hydrodynamic size of PEG-incorporated nanoparticles. c) Transmission Electron Micrographs depict spherical morphology. d) surface charge of nanoparticles after incorporation. e) In vivo nanoparticle administration scheme following pre-injection of hyperosmotic mannitol. f) representative vivo imaging system (IVIS) image of GFP expression in the brain and g) liver following the protocol shown in e).h) GFP expression plotted as the percentage total emission from the brain and liver calculated from ex vivo IVIS fluorescence images of M30 D90.Data represented as mean ± SD, n=3.

In vivo transfection was qualitatively assessed via ex vivo fluorescence imaging. The experimental procedure involved intraperitoneal injection of hyperosmotic mannitol, followed by intravenous administration of nanoparticles at a dose of 0.3mg/kg. After 24 hours, mice were euthanized, and organs were isolated for imaging with an in vivo imaging system (IVIS) (Figure 5e). Strong GFP fluorescence was evident in the brain with M30 D90 including some signal in the liver. Transfection was only observed in the liver with D90 (Figures 5f and 5g). No signals were detected in the lung or spleen for D90 or M30 D90 (Figure S6). Notably, in the case of M30 D90, approximately 10% of the total signal was from the brain (Figure 5h). However, M30 D90 administration without pre-mannitol injection did not result in transfection in the brain (Figure S7a). Instead, transfection was only observed in the liver (Figure S7b). We also noted a decrease in signal in the liver in this case as compared to the scenario where M30 D90 was administered after mannitol pre-injection, suggesting systemic mannitol may enhance overall transfection due to caveolae induction (Figure S7c). Since pre-mannitol injection did not result in brain transfection with D90, and without pre-mannitol injection, M30 D90 did not transfect the brain, both caveolae induction and the caveolae selectivity of nanoparticles were essential for brain delivery.

### Uptake, localization, and safety profile of M30 D90 nanoparticles after in vivo administration

To further confirm the penetration of M30 D90 across BBB, we checked the brain uptake of nanoparticles formed with ATTO647N labeled pDNA. Nanoparticles were administered intravenously following intraperitoneal mannitol injection and after 6 hours, mice brain was isolated, fixed, and sectioned for fluorescence imaging. Punctas of nanoparticles were observed in the brain section confirming delivery across the BBB (Figure S8). To examine the uptake of M30 D90 across various cell types, we conducted colocalization experiments by immunostaining with cell type-specific markers for neurons (MAP2), microglia (Iba-1), and astrocytes (GFAP). Our findings revealed colocalization of M30 D90 with neurons, as evidenced by overlapping signals with MAP2 staining (Figure 6a). We did not observe significant colocalization with microglia (Iba-1) or astrocytes (GFAP), suggesting a potential preference for uptake in neurons over other cell types (Figure 6b and 6c).

**Figure 6:**
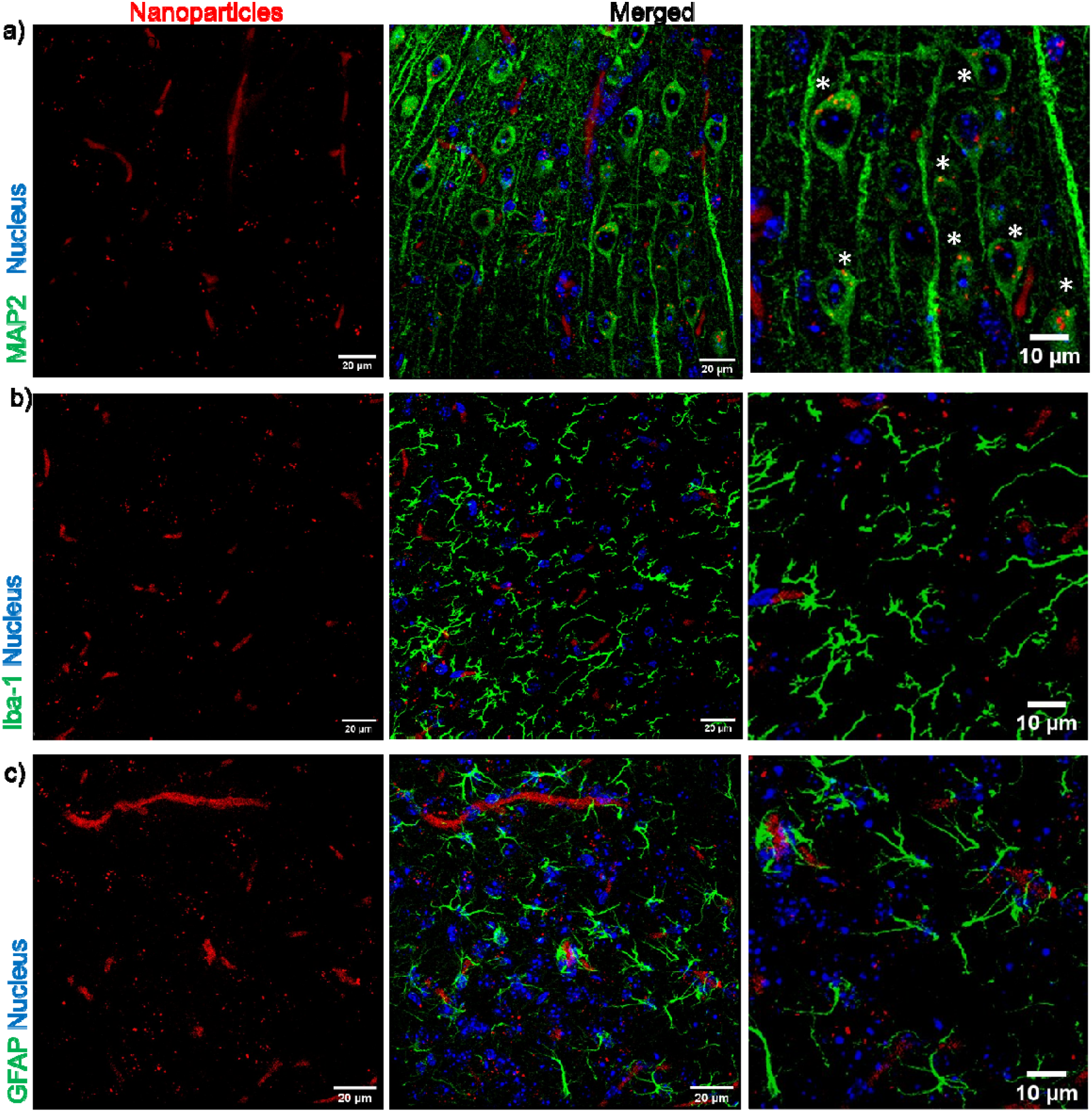
M30 D90 uptake is selective to neurons in the brain in vivo. The biodistribution of M30 D90 in vivo was analyzed by intravenous administration of ATT0647N labeled pDNA following caveolae induction with mannitol administered via intraperitoneal injection. Immunohistochemistry with markers indicates colocalization of M30 D90 nanoparticles (red), a) with neurons (MAP-2, green), b) but not with microglia (Iba-1, green), or c) astrocytes (GFAP, green). Asterisks indicate colocalization in neurons. Z-stacked images depict cell markers (green), nanoparticles (red), and nuclei (blue). Scale bar:20μm. Zoomed-in images scale bar:10μm.

We performed liver enzyme assays to check any hepatotoxicity with M30 D90. No significant difference was noted in the levels of alanine transaminase (ALT) (Figure S9a) and aspartate transaminase (AST) (Figure S9b) between the control and M30 D90. Next, we assessed brain histology one day post-M30 D90 nanoparticle exposure. No necrosis or inflammation was observed with H&E-stained brain sections (Figure S9c).

## Discussion

Most strategies for brain delivery focus on increasing nanoparticle transcytosis with receptor-ligand engagement. Despite success with the delivery of antibodies and oligonucleotides^[44,45]^, these strategies could not be adopted effectively for nanoparticle systems carrying macromolecular cargoes like plasmids. The observation of a physiological shift from clathrin to caveolae-mediated transcytosis in age and some disease states has implied the utilization of caveolae modulation at the BBB as a way to overcome BBB^[10,19]^. Consequently, strategies that modulate caveolae-mediated transcytosis directly or indirectly at the BBB are emerging^[24,25,46,47]^. These studies promote transiently upregulating caveolae-mediated transcytosis at the BBB to transport small and large cargoes to the brain. It is important to note that the overall permeability increases with caveolae modulators, which might not specifically increase the permeability of the drug or the delivery agent.

Nanoparticle-based delivery systems typically use multiple routes to enter the cell in vitro. One way to enhance selectivity for caveolae-mediated endocytosis is by employing osmotic stimuli.^[33]^ In our previous work, we developed backbone-modified sugar alcohol doped PBAE for caveolae selective entry.^[34]^ The modification was carried out in the DD90-Sc12-E63 backbone identified to target the liver in vivo.^[28]^ To improve liver de-targeting and possible improvement in delivery to the brain, we leveraged the chemical versatility of PBAE polymers documented in the literature. As the diacrylate group, we selected bisphenol A ethoxylate diacrylate (D) over bisphenol A glycerolate (DD) to reduce liver accumulation.^[35]^ We synthesized mannitol-modified polymers in the D90-Sc12-E63 backbone (M30 D90). Mannitol-modified polymers retained nucleic acid binding properties and osmotic properties of mannitol. M30 D90 increased cav-1 levels in brain endothelial cells supporting the involvement and preference for caveolae-mediated endocytosis. The transport of M30 D90 nanoparticles was also better than D90 across brain endothelial cells in a transwell culture providing evidence that targeting caveolar transport might lead to better penetration across the BBB. Moreover, the improved release profile of M30 D90 and selective caveolae-mediated entry likely contributed to a three-fold difference in transfection efficiency compared to D90 in neuronal cells.

While M30 D90 increases cav-1 levels in cultured brain endothelial cells, it is unlikely to be adequate for enabling passage across the BBB in vivo. In a healthy brain, non-specific caveolae-mediated endocytosis is kept low.^[38]^For nanoparticles to utilize this pathway, caveolar transport needs to be induced. Transient upregulation of cav-1 has been reported to increase the transcytosis rate at the BBB. Chemical cues like vascular endothelial growth factor^[48]^ and low-dose methamphetamine^[24]^ have been reported to increase caveolae-mediated transcytosis. Systemic administration of hyperosmolar mannitol has been reported to increase pinocytic vesicles in the brain.^[36,40]^ There have been speculations that these pinocytic vesicles could be caveolae.^[37]^ Earlier reports have noted increased transgene expression with rAAV-mediated delivery followed by systemic mannitol injection.^[41,42]^ With this evidence, we employed hyperosmolar mannitol for caveolae induction at the BBB. When administered intraperitoneally, hyperosmolar mannitol increased cav-1 levels in the brain in under 30 minutes as observed by immunostaining. We show that changes in cav-1 might not be affecting paracellular permeability as observed with the state of tight junction marker (occludin-1). We suspect vasodilation with slightly thicker blood vessels observed in the images with immunostaining of occludin-1. Our results indicate that hyperosmolar mannitol administered systemically can be used for caveolae induction without affecting tight junction integrity. Increased intracellular permeability without paracellular permeability changes have been reported with age.^[49]^

PEG-incorporated M30 D90 crossed the BBB to transfect the brain following caveolae induction with hyperosmolar mannitol at a low dose of 0.3mg/kg of pDNA. Without mannitol pre-injection, M30 D90 did not demonstrate transfection in the brain. On the other hand, even with pre-mannitol injection, unmodified polymer-D90 also failed to show transfection in the brain. These findings highlight the importance of caveolae induction at the BBB and nanoparticle selectivity for delivery through caveolae-mediated transcytosis. To check if the mannitol modification is helpful even with a different diacrylate, we also checked the transfection with M30 DD90 (Mannitol modified in DD90-Sc-12 backbone) which we have reported earlier. We found that M30 DD90 also transfected the brain in the caveolae induction model of systemic mannitol. However, the levels of transfection in the brain were less and the brain-to-liver ratio was lower than that observed with M30 D90 (data not included). These results strongly indicate that the mannitol modification needs to be coupled with appropriate monomer composition and combined with pre-mannitol injection to achieve efficient transfection in the brain. M30 D90 uptake was predominantly found in neurons making it suitable for cell type-specific applications. However, the reason for the specificity of M30 D90 to neurons remains elusive, it warrants further exploration of the monomer compositions in play.

In summary, this study highlights the role of caveolae targeting for delivering nucleic acids to the brain. We strategically designed a nanoparticle system based on PBAE polymer modified with mannitol to target the caveolae-mediated endocytosis pathway for crossing the BBB. We also used systemic mannitol injection to enhance caveolae formation at the BBB in order to facilitate the entry of these nanoparticles. We establish that this combination is critical for the delivery. We also establish that the monomer composition is important for liver de-targeting. Although we observe higher localization of these nanoparticles in the neurons, it is possible that targeted caveolae induction with low-dose focused ultrasound can enable further localized access to specific brain regions. We believe this provides the foundation for exploring this system for therapeutic genes to treat CNS disorders.

## Materials and Methods

Bisphenol A ethoxylate diacrylate (413550), 4-(2-aminoethyl) morpholine (S90; CAS 2038-031), 1-Dodecylamine (Sc12; CAS 124-22-1), Diethylenetriamine (E63; CAS 111400), Mannitol (CAS 69658), Pyridine (CAS 110861), Chlorpromazine Hydrochloride (C0982), Nystatin (N9150), Cytochalasin D (C8273), ATTO488 amine (74417), ATTO647N amine (95349), SPB (succinimidyl-[4-(psoralen-8-yloxy)] butyrate) (803545), DMSO-D6, Dimethylformamide (CAS 68-12-2), Diethyl ether (CAS 60-29-7), Thiazolyl Blue Tetrazolium Bromide (M5655), DMG PEG 2000 (880151P-1G), Spectra-Por Float-A-Lyser G2 (Z726710-12EA), ALT activity assay (MAK052), AST activity assay (MAK055)were purchased from Sigma-Aldrich. Acryloyl chloride (CAS 2123990) was purchased from Alfa Aesar. The pEGFP-C1 plasmid (4.7 kb) was amplified in Escherichia coli DH-5α and pDNA isolated with GenEluteTM Sigma-Aldrich. The primary antibody against Caveolin-1 (ab2910), GFAP (ab7260), MAP-2(ab32454), and Iba-1 (ab178846) was purchased from Abcam. Occludin (3E11.11) was procured from DSHB. Secondary antibodies Alexa Fluor 594 goat anti-rabbit (A11012), Alexa fluor 488 goat anti-rabbit (A11008), Alexa Fluor 568 goat anti-rat (A11077), and Prolong Gold AntiFade with DAPI (P3693) were procured from Thermo Fisher Scientific. Costar6.5mm transwell permeable support with 3μm pore polyester membrane (3472) was procured from corning.

### Mannitol-modified polymer synthesis and characterization

The polymers were synthesized as previously reported^[34]^. Briefly, diacrylate - Bisphenol A ethoxylate diacrylate (D), hydrophilic amine 4-(2-aminoethyl) morpholine (S90), and hydrophobic amine – (1-Dodecylamine (Sc-12) were reacted for 24 hours in DMF to synthesize the base polymer of D90. For mannitol modification, mannitol diacrylate was substituted at a 30% molar ratio to diacrylate – Dtofrom M30 D90. Both the polymers were capped with Diethylenetriamine (E63) in THF for 2 hours toget the final polymers. Polymers were purified with diethyl ether and vacuum-dried. Polymers were characterized with 400 MHz Bruker (400 MHz for 1H-NMR)to confirm mannitol incorporation in the backbone. The molecular weight of the polymers was evaluated with GPC (Malvern Omnisec instrument having an RI detector and Shodex KD-806 M).The polymers were dissolved in anhydrous DMSO and stored at −20°C until further use.

### Nucleic acid binding assay

Polymers were evaluated for nucleic acid binding with RiboGreen. D90 and M30 D90 polymers were serially diluted from 100μg/ml concentration in 25mM sodium acetate buffer. The pDNA (pEGFP-C1) solution was prepared at a 1μg/mL stock concentration with RiboGreen dye in 25 mM sodium acetate buffer. 25 μL of the polymer solution was mixed with 75μL of nucleic acid/RiboGreen solution in 96 well-black bottom plates. The samples were incubated for 20 minutes at 37°C and fluorescent reading was taken in Tecan multi-plate reader excited at 490nm the fluorescence emission intensity was measured at 530nm. IC50 value was obtained by plotting the fluorescence quenching as a function of polymer concentration and fitting a sigmoid curve to the data. Lower IC50 values indicate higher binding and vice versa.

### Osmolarity measurement

The osmolarity of the polymers was measured at 2.5 and 5 % of polymer concentration in 100 mM sodium acetate buffer at pH 5.2. Reading was measured using a cryoscopic osmometer (OSMOMAT 3000 Microprocessor).

### Nanoparticle formation and characterization

Nanoparticles were formed by pipette mixing pDNA and polymer at 1:60 (w/w) in 25mM sodium acetate (pH 5.2). For size measurements, 4μg of plasmid was used in nanoparticle formation with polymers. The nanoparticles were diluted in MilliQ at 0.002μg/μl concentration and for Zeta potential measurement nanoparticles were diluted in 10mM NaCl and analyzed using Zetasizer Nano ZS (Malvern Instruments, UK).

### Gel electrophoresis

Complexation and release studies were done using agarose gel electrophoresis. Agarose gels were cast with 1% agarose dissolved in Tris-Acetate-EDTA buffer. 25ng of pDNA was loaded per well. To pDNA different w/w ratios of polymer was added (1:0.5,1:1,1:5,1:10,1:201:30,1:40,1:60). For release studies, the pDNA to polymer ratio was fixed to a 1:60 (w/w) ratio and the heparin weights were set in gradients 1:0.25, 1:0.5, 1:1, 1:2,1:4, 1:8, 1:16, 1:36 to the polymers. Orange loading dye was added to the sample before loading the sample to the gel, electrophoresis was run for 100V for 10 minutes and the gel was visualized using gel doc.

### Transmission Electron Microscopy

TEM imaging was conducted to gain insight into the structure and localization of DNA within the nanoparticle. Assembled nanoparticles were placed onto 200 mesh copper grids. Subsequently, the copper grids were washed using ultrapure water to eliminate buffer salts. Afterward, the grids were rinsed with 1% uranyl acetate in ultrapure water and air-dried overnight.The samples were then analyzed using the TECNAI G2 20 Twin electron microscope, capturing 10–15 fields for each sample.

### Cell culture

SHSY5Y cell line (Human neuroblastoma) was provided by Dr. Beena Pillai at CSIR-IGIB. bEND.3 cell line (Mouse brain microvascular endothelium) was procured from AddexBio. Cells were grown in DMEM media from Gibco supplemented with 10% fetal bovine serum.

### Plasmid labelling

DNA was labeled with ATTO488 and ATTO647N as reported earlier^[50]^. 100μl (1μg/μL) of pDNA was incubated with 12.5μL (1μg/μL) of NHS psoralen in a round bottom 96-well plate for crosslinking under a 365nm UV lamp for 25 minutes. Following crosslinking reaction ATTO488 or ATTO647N amine was added and incubated for 1 hour at room temperature. The labeled plasmid was purified by ethanol precipitation. pDNA was resuspended in nuclease-free water and stored at −20°C for further use.

### Mechanism of endocytosis

The mechanism of endocytosis was assayed in bEND.3 and SHSY5Y cell lines. Cells were seeded at a density of 75,000 cells per 24-well plate. Confluent wells were treated with endocytosis inhibitors at the following concentrations, Chlorpromazine (10 μg/mL),Cytochalasin D (500nM), and Nystatin (50μg/ml). Following 1 hour incubation with the inhibitors, nanoparticles of D90, and M30 D90 prepared with ATTO488 labeled pDNA were added, and the cells were processed for flow cytometry after 2 hours. Data was collected on a BD Accuri C6 flow cytometer and analyzed using BD Accuri C6 software.

### Immunocytochemistry in bEND.3

bEND.3 cells were seeded onto coverslips in a 6-well plate, at a density of 100,000 cells per well, allowing them to adhere overnight. Cells were incubated with nanoparticles formed with 2μg pDNA per well for 30 minutes and were subsequently fixed using 4% paraformaldehyde for 15 minutes at room temperature. Following fixation, cells were permeabilized using a permeabilization buffer (0.1% Triton-X100 in PBS) for 5 minutes. After that, the cells were immersed in a blocking buffer (5% BSA in PBS with 0.1% Tween 20) for 1 hour at room temperature. The primary antibody (anti-caveolin 1 antibody), at 1:500 dilution buffer, was added to the blocking buffer and left overnight at 4°C in a humid chamber. Coverslips were washed with wash buffer (0.1% Tween 20 in PBS) and secondary antibody (goat anti-rabbit secondary antibody Alexa Fluor 594), diluted 1:800 in the blocking buffer, added to the coverslip, and incubated for 1 hour. After washing the coverslips 3 times with wash buffer the coverslips were mounted with Prolong antifade withDAPI(Life Technologies). The images were taken with the Life Technologies Floid Cell Imaging Station.

### qRT-PCR

Following 30 minutes of nanoparticle treatment,total messenger RNA was isolated from bEND.3 cells using trizol method with TRI Reagent.Following reverse transcription with the iScript cDNA synthesis kit (Bio-Rad), the transcripts were quantified using KAPA SYBR® FAST according to the manufacturer’s instructions in the PCR instrument (BioRad). Fold changes in expression were calculated using the ΔΔCT method. The GAPDH gene was used to normalize results. The primer sequences were taken from the previous report and are listed in supplementary Table S3^[20]^.

### Penetration assay

bEnd.3 cells were seeded on the upper surface of the membrane in polyester transwell inserts (3μm pore size, and 6.5mm diameter) at a density of 25,000 cells per well. Media were changed every other day. Cells were cultured for 5-7 days until a confluent monolayer was formed. Nanoparticles formed with ATTO488 labeled plasmid (at a dose of 500ng) were added to the upper chamber in flurobrite media containing FBS. The penetration of nanoparticles across the cells was assessed by measuring the nanoparticle concentration from the basal well media after 5 hours of incubation. Nanoparticle concentration was determined by NTA (NanoSight NS300), samples were loaded on the top plate and analyzed through a long pass filter with a wavelength cut-off of 500nm to identify the nanoparticles. For each sample, 5 × 60s videos were recorded at 25°C and the concentrations were obtained using NanoSight software.

### Transfection, and cytotoxicity

SHSY5Y cells were seeded at 50,000 cells per well in 24-well plates. Nanoparticles formed with GFP reporter plasmid were treated to the cells in complete media for 24 hours, with 300ng pDNA per 24-well plate. After 24 hours, cytotoxicity was estimated by MTT assay and transfection efficiency was measured through flow cytometry. Fluorescence images of transfection were taken using the Life Technologies Floid Cell Imaging Station.

### Nanoparticle assembly for in vivo delivery

PEG lipid was introduced in the nanoparticle system to enable systemic stability in serum for in vivo delivery. Nanoparticles were formulated with a 1:60 w/w ratio. PEG lipid (10% by weight of polymer) and polymer in absolute ethanol were mixed with DNA in sodium acetate buffer (pH 5.2). The pDNA and polymer-PEG lipid solution was mixed by pipetting at a 3:1 volume ratio and incubated for 10 minutes at room temperature for nanoparticle formation. The nanoparticles were dialyzed for 2 hours with a 20kDa Float-A-Lyser G2 dialysis device against PBS at 4°C.

### Animal studies

The Institutional Animal Ethics Committee approved all animal procedures. All experiments were performed in 6 to 8-week-old Balb/C female mice.

### Caveolae induction

For caveola induction, 20% mannitol solution prepared in saline was injected intraperitoneally (0.8M mannitol). To check the effect of systemic mannitol in caveolar transport, mice brain was collected after 30 minutes. Brain tissue was fixed in 4% (w/v) PFA overnight at 4°C before preservation in 30% (w/v) sucrose in PBS. The brain was embedded in OCT compound (Leica) and sectioned at a thickness of 15⍰μm on a freezing–sliding microtome. Sections were blocked with 5% horse serum and 1% bovine serum albumin followed by the addition of primary antibodies (Caveolin-1 and Occludin-1) for overnight incubation at 4°C. After washing steps, secondary antibodies were added according to the manufacturer’s dilution. The tissue samples were sealed with a coverslip with antifade mountant containing DAPI. Images were acquired using Leica SP8 confocal microscope.

### In vivo transfection

In vivo transfection was checked with GFP reporter plasmid (pEGFP-C1). Before administration of nanoparticles caveolae were induced with mannitol as mentioned earlier. After 5 minutes of mannitol injection administered intraperitoneally, PEG-incorporated nanoparticles encapsulating reporter plasmid were injected via the lateral tail vein at a 0.3mg/kg dose. After 24 hours, animals were euthanized via thiopental injection, and selected organs were extracted for imaging with IVIS (Spectral Instruments Imaging System Lago X). For transfection without caveolae induction, only M30 D90 was administered intravenously and organ fluorescence was measured with IVIS after 24 hours.

### In vivo uptake

M30 D90 uptake in the brain was checked with nanoparticles formed with ATTO647N labeled plasmid. Following caveolae induction with hyperosmolar mannitol,a 0.3mg/kg dose of ATTO647N labeled M30 D90 nanoparticles was administered intravenously. After 6 hours,the brain was isolated, fixed, and sectioned for immunostaining. Sections were blocked with 5% horse serum before incubation at 4°C with primary antibodies MAP2 (neuronal marker), GFAP (astrocyte marker), or Iba-1 (microglial marker)(dilutions were as per manufacturer’s protocol). Sections were washed, stained with Alexa Fluor-conjugated secondary antibodies, and mounted with antifade and coverslip before imaging on a Leica SP8 confocal microscope.

### In vivo safety

For in vivo safety profiling, blood was collected after 24 hours of nanoparticle administration. Blood was centrifuged at 13,000 rpm for 10 minutes to retrieve serum. Liver enzyme assays AST and ALT activity were measured according to manufacturers’ protocol (Sigma-Aldrich, St. Louis, MO). Brain sections were stained for H & E and imaged on Nikon digital sight DS-U3 under 4X magnification.

### Statistics

Statistical analyses were performed using GraphPad Prism software (GraphPad Software, Inc). Results are depicted as the mean and standard with three repeats if not stated otherwise. Significance was calculated with two-way ANOVA. Statistical significance was denoted as follows: *p < 0.05, **p < 0.01, ***p < 0.001, ****p < 0.0001. All experiments were done in triplicates until and otherwise mentioned.

## DATA AND CODE AVAILABILITY

The data presented in this study are available on request from the corresponding authors.

## ACKNOWLEGEMENTS

We thank the Council of Scientific and Industrial Research (MLP2011) for the financial support to carry out the work. CSIR-GATE supports B.R.G. for the fellowship. We thank Koushikafor helping with confocal and the National Institute of Immunology for the technical support with TEM, tissue sectioning, and H&E staining. Figures were created with Adobe Illustrator and bio render.

## AUTHOR CONTRIBUTIONS

B.R.G.: Conceptualization, methodology, Formal analysis, investigation, visualization, writing – original draft preparation, writing – review & editing. C.M.: Formal analysis, investigation. A.K.: Formal analysis, investigation. S.G.: Project administration, supervision. A.P.:Project administration, supervision, writing – review & editing. M.G.: conceptualization, funding acquisition, investigation, project administration, supervision, writing – review & editing

## Reference

[1] M. Zheng, W. Tao, Y. Zou, O. C. Farokhzad, B. Shi, Trends Biotechnol 2018, 36, 562–575.

[2] Z. G. Lu, J. Shen, J. Yang, J. W. Wang, R. C. Zhao, T. L. Zhang, J. Guo, X. Zhang, Signal Transduct Target Ther 2023, 8, DOI 10.1038/s41392-022-01298-z.

[3] B. W. Chow, C. Gu, Trends Neurosci 2015, 38, 598–608.

[4] W. A. Banks, Nat Rev Drug Discov 2016, 15, 275–292.

[5] G. C. Terstappen, A. H. Meyer, R. D. Bell, W. Zhang, Nat Rev Drug Discov 2021, 20, 362–383.

[6] M. Sela, M. Poley, P. Mora-Raimundo, S. Kagan, A. Avital, M. Kaduri, G. Chen, O. Adir, A. Rozencweig, Y. Weiss, O. Sade, Y. Leichtmann-Bardoogo, L. Simchi, S. Aga-Mizrachi, B. Bell, Y. Yeretz-Peretz, A. Z. Or, A. Choudhary, I. Rosh, D. Cordeiro, S. Cohen-Adiv, Y. Berdichevsky, A. Odeh, J. Shklover, J. Shainsky-Roitman, J. E. Schroeder, D. Hershkovitz, P. Hasson, A. Ashkenazi, S. Stern, T. Laviv, A. Ben-Zvi, A. Avital, U. Ashery, B. M. Maoz, A. Schroeder, Advanced Materials 2023, 35, DOI 10.1002/adma.202304654.

[7] E. A. Wyatt, M. E. Davis, Mol Pharm 2020, 17, 717–721.

[8] L. E. Oikari, R. Pandit, R. Stewart, C. Cuní-López, H. Quek, R. Sutharsan, L. M. Rantanen, M. Oksanen, S. Lehtonen, C. M. de Boer, J. M. Polo, J. Götz, J. Koistinaho, A. R. White, Stem Cell Reports 2020, 14, 924–939.

[9] G. C. Tracy, K. Y. Huang, Y. T. Hong, S. Ding, H. A. Noblet, K. H. Lim, E. C. Kim, H. J. Chung, H. Kong, Nano Lett 2023, 23, 10971–10982.

[10] A. C. Yang, M. Y. Stevens, M. B. Chen, D. P. Lee, D. Stähli, D. Gate, K. Contrepois, W. Chen, T. Iram, L. Zhang, R. T. Vest, A. Chaney, B. Lehallier, N. Olsson, H. du Bois, R. Hsieh, H. C. Cropper, D. Berdnik, L. Li, E. Y. Wang, G. M. Traber, C. R. Bertozzi, J. Luo, M. P. Snyder, J. E. Elias, S. R. Quake, M. L. James, T. Wyss-Coray, Nature 2020, 583, 425–430.

[11] K. Tachibana, Y. Hashimoto, K. Shirakura, Y. Okada, R. Hirayama, Y. Iwashita, I. Nishino, Y. Ago, H. Takeda, H. Kuniyasu, M. Kondoh, Journal of Controlled Release 2021, 336, 105–111.

[12] N. H. On, P. Kiptoo, T. J. Siahaan, D. W. Miller, Mol Pharm 2014, 11, 974–981.

[13] B. M. D. C. Godinho, N. Henninger, J. Bouley, J. F. Alterman, R. A. Haraszti, J. W. Gilbert, E. Sapp, A. H. Coles, A. Biscans, M. Nikan, D. Echeverria, M. DiFiglia, N. Aronin, A. Khvorova, Molecular Therapy 2018, 26, 2580–2591.

[14] A. M. Sonabend, A. Gould, C. Amidei, R. Ward, K. A. Schmidt, D. Y. Zhang, C. Gomez, J. F. Bebawy, B. P. Liu, G. Bouchoux, C. Desseaux, I. B. Helenowski, R. V. Lukas, K. Dixit, P. Kumthekar, V. A. Arrieta, M. S. Lesniak, A. Carpentier, H. Zhang, M. Muzzio, M. Canney, R. Stupp, Lancet Oncol 2023, 24, 509–522.

[15] Y. H. Lao, R. Ji, J. K. Zhou, K. J. Snow, N. Kwon, E. Saville, S. He, S. Chauhan, C. W. Chi, M. S. Datta, H. Zhang, C. H. Quek, S. S. Cai, M. Li, Y. Gaitan, L. Bechtel, S. Y. Wu, C. M. Lutz, R. Tomer, S. A. Murray, A. Chavez, E. E. Konofagou, K. W. Leong, Proc Natl Acad Sci U S A 2023, 120, DOI 10.1073/pnas.2302910120.

[16] R. H. Kofoed, C. L. Dibia, K. Noseworthy, K. Xhima, N. Vacaresse, K. Hynynen, I. Aubert, Journal of Controlled Release 2022, 351, 667–680.

[17] G. Kwak, A. Grewal, H. Slika, G. Mess, H. Li, M. Kwatra, A. Poulopoulos, G. F. Woodworth, C. G. Eberhart, H. Ko, A. Manbachi, J. Caplan, R. J. Price, B. Tyler, J. S. Suk, ACS Nano 2024, DOI 10.1021/acsnano.4c05270.

[18] Y. L. Zhao, J. N. Song, M. Zhang, Rev Neurosci 2014, 25, 247–254.

[19] A. G. Sorets, J. C. Rosch, C. L. Duvall, E. S. Lippmann, Curr Opin Chem Eng 2020, 30, 86–95.

[20] D. E. Tylawsky, H. Kiguchi, J. Vaynshteyn, J. Gerwin, J. Shah, T. Islam, J. A. Boyer, D. R. Boué, M. Snuderl, M. B. Greenblatt, Y. Shamay, G. P. Raju, D. A. Heller, Nat Mater 2023, 22, 391–399.

[21] S. Lei, J. Li, J. Yu, F. Li, Y. Pan, X. Chen, C. Ma, W. Zhao, X. Tang, Int J Oral Sci 2023, 15, DOI 10.1038/s41368-022-00215-y.

[22] H. Salimi, M. D. Cain, X. Jiang, R. A. Roth, W. L. Beatty, C. Sun, W. B. Klimstra, J. Hou, R. S. Klein, 2020, DOI 10.1128/mBio.

[23] B. J. Andreone, B. W. Chow, A. Tata, B. Lacoste, A. Ben-Zvi, K. Bullock, A. A. Deik, D. D. Ginty, C. B. Clish, C. Gu, Neuron 2017, 94, 581-594.e5.

[24] J. H. Chang, C. Greene, K. Frudd, L. Araujo dos Santos, C. Futter, B. J. Nichols, M. Campbell, P. Turowski, Cell Rep Med 2022, 3, DOI 10.1016/j.xcrm.2021.100497.

[25] R. Pandit, W. K. Koh, R. K. P. Sullivan, T. Palliyaguru, R. G. Parton, J. Götz, Journal of Controlled Release 2020, 327, 667–675.

[26] Z. G. Lu, J. Shen, J. Yang, J. W. Wang, R. C. Zhao, T. L. Zhang, J. Guo, X. Zhang, Signal Transduct Target Ther 2023, 8, DOI 10.1038/s41392-022-01298-z.

[27] L. Rotolo, D. Vanover, N. C. Bruno, H. E. Peck, C. Zurla, J. Murray, R. K. Noel, L. O’Farrell, M. Araínga, N. Orr-Burks, J. Y. Joo, L. C. S. Chaves, Y. Jung, J. Beyersdorf, S. Gumber, R. Guerrero-Ferreira, S. Cornejo, M. Thoresen, A. K. Olivier, K. M. Kuo, J. C. Gumbart, A. R. Woolums, F. Villinger, E. R. Lafontaine, R. J. Hogan, M. G. Finn, P. J. Santangelo, Nat Mater 2023, 22, 369–379.

[28] Y. Rui, D. R. Wilson, S. Y. Tzeng, H. M. Yamagata, D. Sudhakar, M. Conge, C. A. Berlinicke, D. J. Zack, A. Tuesca, J. J. Green, H E A L T H A N D M E D I C I N E High-Throughput and High-Content Bioassay Enables Tuning of Polyester Nanoparticles for Cellular Uptake, Endosomal Escape, and Systemic in Vivo Delivery of MRNA, 2022.

[29] A. F. Rodrigues, C. Rebelo, S. Simões, C. Paulo, S. Pinho, V. Francisco, L. Ferreira, Advanced Science 2023, 10, DOI 10.1002/advs.202205475.

[30] A. Mangraviti, S. Y. Tzeng, K. L. Kozielski, Y. Wang, Y. Jin, D. Gullotti, M. Pedone, N. Buaron, A. Liu, D. R. Wilson, S. K. Hansen, F. J. Rodriguez, G. D. Gao, F. Dimeco, H. Brem, A. Olivi, B. Tyler, J. J. Green, ACS Nano 2015, 9, 1236–1249.

[31] L. Gao, C. Shi, Z. Yang, W. Jing, M. Han, J. Zhang, C. Zhang, C. Tang, Y. Dong, Y. Liu, C. Chen, X. Jiang, J Nanobiotechnology 2023, 21, DOI 10.1186/s12951-023-01810-9.

[32] P. Mastorakos, C. Zhang, E. Song, Y. E. Kim, H. W. Park, S. Berry, W. K. Choi, J. Hanes, J. S. Suk, Journal of Controlled Release 2017, 262, 37–46.

[33] M. Ariful Islam, T. E. Park, J. Firdous, H. S. Li, Z. Jimenez, M. Lim, J. W. Choi, C. H. Yun, C. S. Cho, Essential Cues of Engineered Polymeric Materials Regulating Gene Transfer Pathways, 2022.

[34] B. Reshma G, C. Miglani, A. Pal, M. Ganguli, Nanoscale 2024, DOI 10.1039/d3nr05300h.

[35] J. C. Kaczmarek, A. K. Patel, L. H. Rhym, U. C. Palmiero, B. Bhat, M. W. Heartlein, F. DeRosa, D. G. Anderson, Biomaterials 2021, 275, DOI 10.1016/j.biomaterials.2021.120966.

[36] A. Hirano, T. Kawanami, J. F. Llena, Electron Microscopy of the Blood-Brain Barrier in Disease, 1994.

[37] S. R. Burks, C. N. Kersch, J. A. Witko, M. A. Pagel, M. Sundby, L. L. Muldoon, E. A. Neuwelt, J. A. Frank, D. M. Holtzman, n.d., DOI 10.1073/pnas.2021915118/-/DCSupplemental.

[38] S. Ayloo, C. Gu, Curr Opin Neurobiol 2019, 57, 32–38.

[39] R. Villaseñor, J. Lampe, M. Schwaninger, L. Collin, Cellular and Molecular Life Sciences 2019, 76, 1081–1092.

[40] T. O. Lilius, K. N. Mortensen, C. Deville, T. J. Lohela, F. F. Stæger, B. Sigurdsson, E. M. Fiordaliso, M. Rosenholm, C. Kamphuis, F. J. Beekman, A. I. Jensen, M. Nedergaard, Journal of Controlled Release 2023, 355, 135–148.

[41] C. Burger, F. N. Nguyen, J. Deng, R. J. Mandel, Molecular Therapy 2005, 11, 327–331.

[42] D. M. McCarty, J. DiRosario, K. Gulaid, J. Muenzer, H. Fu, Gene Ther 2009, 16, 1340–1352.

[43] A. A. Eltoukhy, D. Chen, C. A. Alabi, R. Langer, D. G. Anderson, Advanced Materials 2013, 25, 1487–1493.

[44] S. J. Barker, M. B. Thayer, C. Kim, D. Tatarakis, M. Simon, R. L. Dial, L. Nilewski, R. C. Wells, Y. Zhou, M. Afetian, K. S. Chew, J. Chow, A. Clemens, C. B. Discenza, J. Dugas, C. Dwyer, T. Earr, C. Ha, D. Huynh, S. Jayaraman, W. Kwan, C. Mahon, M. Pizzo, E. Roche, L. Sanders, A. Stergioulis, R. Tong, H. Tran, J. Zuchero, A. A. Estrada, K. Gadkar, C. M. Koth, P. E. Sanchez, R. G. Thorne, R. J. Watts, T. Sandmann, L. Kane, M. S. Dennis, J. W. Lewcock, S. L. DeVos, n.d., DOI 10.1101/2023.04.25.538145.

[45] T. Okuyama, Y. Eto, N. Sakai, K. Nakamura, T. Yamamoto, M. Yamaoka, T. Ikeda, S. So, K. Tanizawa, H. Sonoda, Y. Sato, Molecular Therapy 2021, 29, 671–679.

[46] Y. Gu, R. Cai, C. Zhang, Y. Xue, Y. Pan, J. Wang, Z. Zhang, FASEB Journal 2019, 33, 441–454.

[47] X. Ju, T. Miao, H. Chen, J. Ni, L. Han, Adv Healthc Mater 2021, 10, DOI 10.1002/adhm.202001997.

[48] L. N. Zhao, Z. H. Yang, Y. H. Liu, H. Q. Ying, H. Zhang, Y. X. Xue, Journal of Molecular Neuroscience 2011, 44, 122–129.

[49] M. J. Cummins, E. T. Cresswell, D. W. Smith, D. Smith, n.d., DOI 10.1101/2024.02.12.580035.

[50] D. R. Wilson, A. Mosenia, M. P. Suprenant, R. Upadhya, D. Routkevitch, R. A. Meyer, A. Quinones-Hinojosa, J. J. Green, J Biomed Mater Res A 2017, 105, 1813–1825.

