## Supplemental for "A combination of systemic mannitol administration and mannitol-modified polyester nanoparticles facilitate gene delivery to the brain through caveolae-mediated endocytosis"

**Table S1** Polymer chemical and molar composition

| Polymer | D | S90 | Sc12 | MDA |
|---|---|---|---|---|
| D90 | 1.2 | 0.5 | 0.5 | |
| M30 D90 | 0.84 | 0.5 | 0.5 | 0.36 |

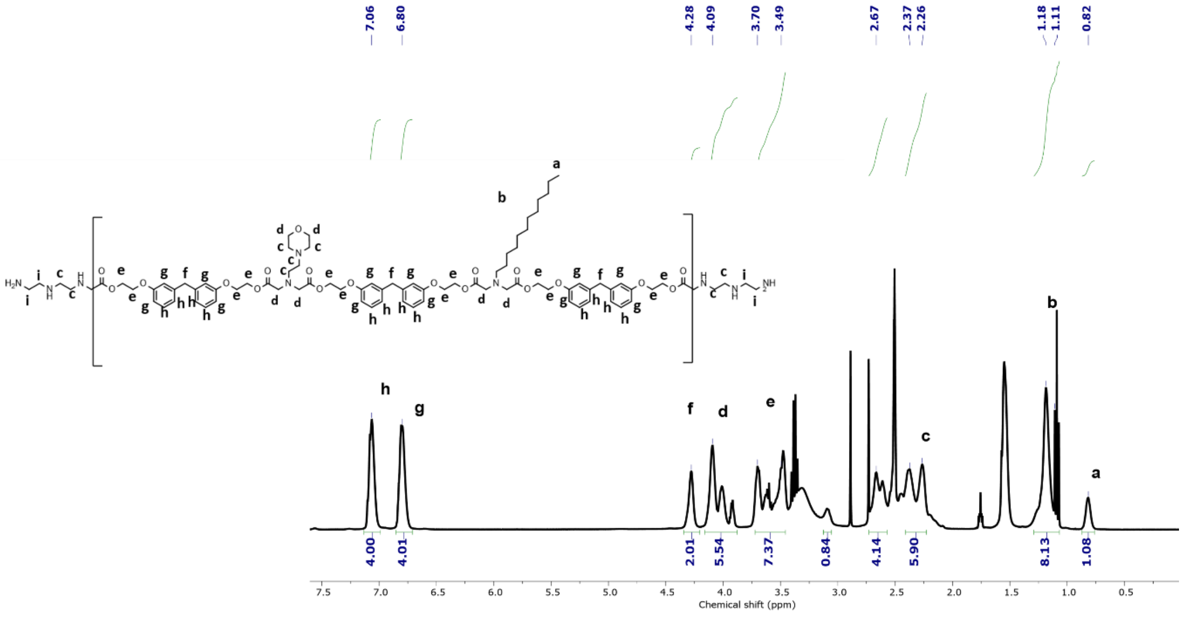

**Figure S1.** $^{1}$H-NMR spectra of D90

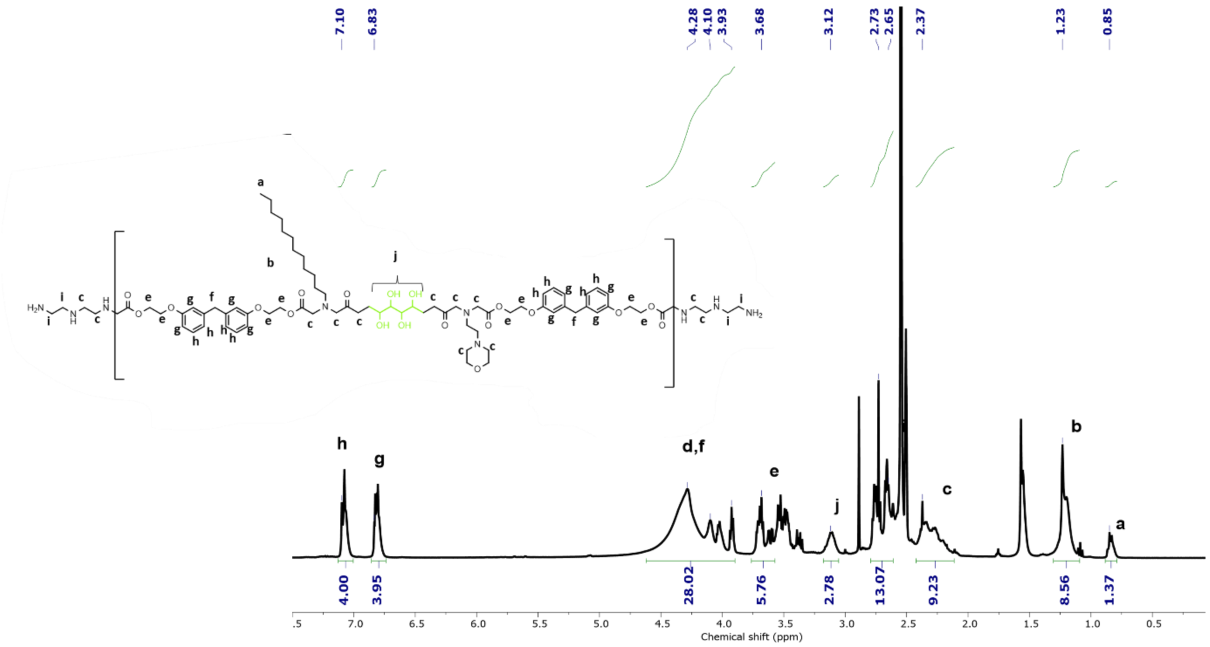


**Figure S2.** ¹H-NMR spectra of M30 D90. The total number of protons in the aromatic ring of the polymer ($a$) = 8. Total no of protons of MDA from ¹H-NMR spectra( $x$ ) = 2.78. Total number of mannitol proton = 8

$$\% \text{ MDA to Polymer} = \frac{\frac{x}{8}}{\frac{x}{8}+\frac{a}{8}} * 100 = \frac{\frac{2.78}{8}}{\frac{2.78}{8}+\frac{8}{8}} * 100 = 26\%$$

**Table S2**. GPC analysis of modified PBAE

| **Polymer** | **Mw** | **Mn** | **PDI** |
|---|---|---|---|
| D90 | 5082 | 6597 | 1.3 |
| M30 D90 | 5901 | 6731 | 1.1 |

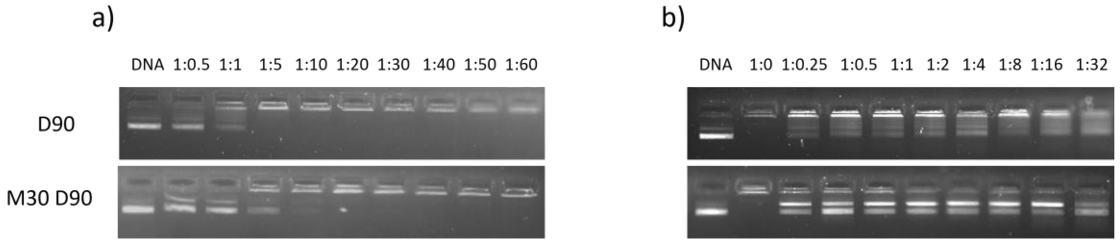


**Figure S3.** DNA condensation and release of polymers. a) D90 and M30 D90 polymers exhibit strong DNA condensation ability and can effectively condense pDNA at the w/w ratios ranging from 1:10 to 1:60. b) When nanoparticles formed with 1:60 w/w (pDNA to polymer) were treated with a heparin gradient, pDNA was released completely in M30 D90 and not in D90.

**Table S3.** qRT-PCR primer sequence

| Primer ID | Sequence |
|---|---|
| mCAV1 FP | GCGACCCCAAGCATCTCA |
| mCAV1 RP | ATGCCGTCGAAACTGTGTGT |
| mGAPDH FP | CGTCCCGTAGACAAAATGGT |
| mGAPDH RP | TCAATGAAGGGGTCGTTGAT |

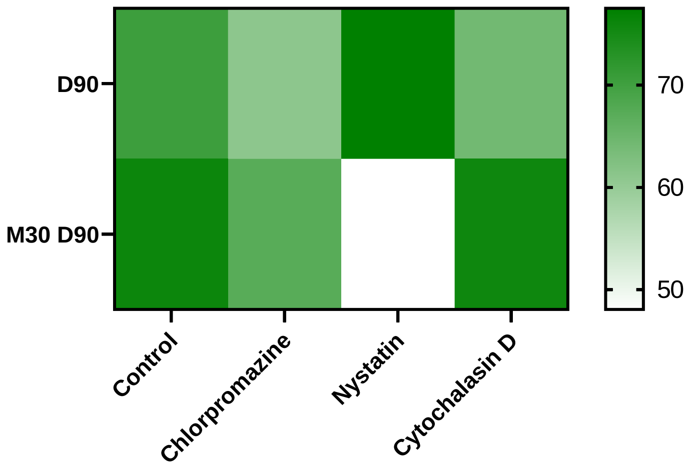


**Figure S4.** Mechanism of endocytosis in SHSY5Y.

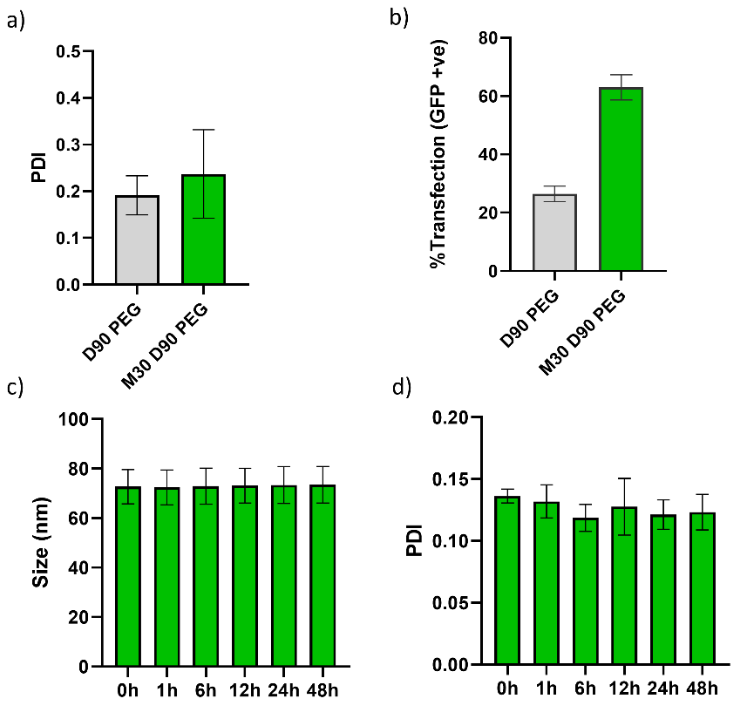

**Figure S5.** PEG-incorporation a) PDI and b) transfection efficiency in SHSY5Y. The stability of PEG incorporated M30 D90 was assessed with c) size and d) PDI as a function of time.

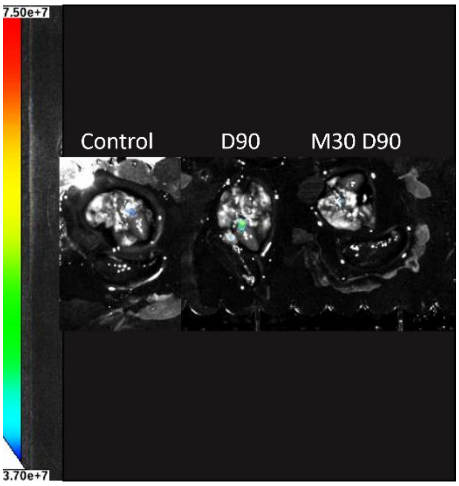


**Figure S6.** Representative IVIS images of GFP expression show no signal in the lung and spleen.

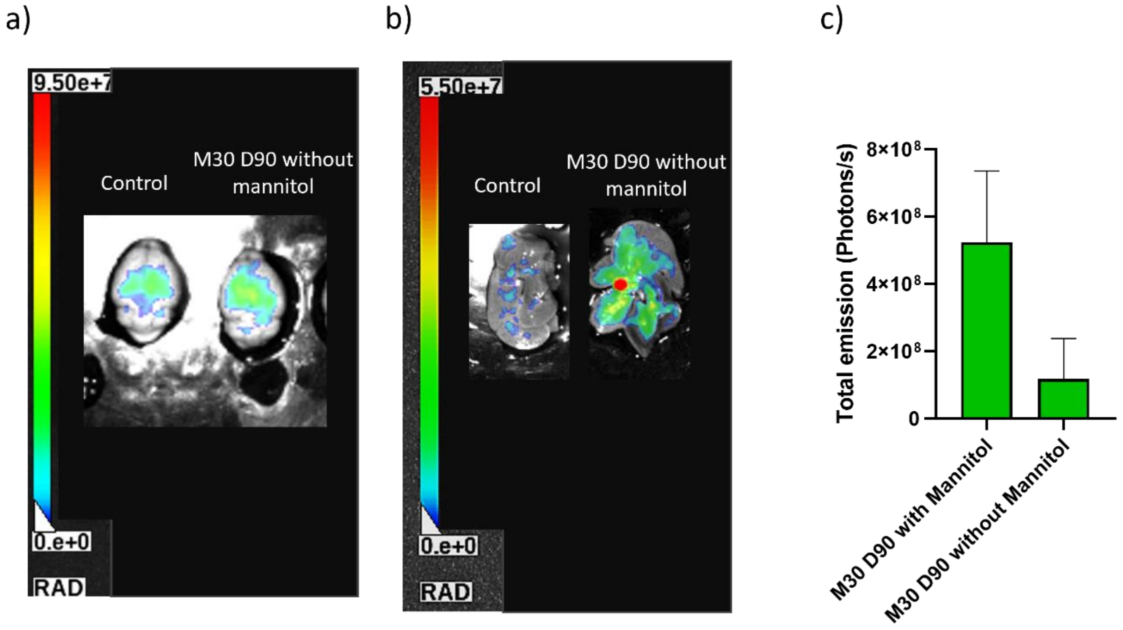


**Figure S7.** M30 D90 formulated with GFP reporter plasmid injected intravenously without pre-mannitol injection (pDNA at a dose of 0.4mg/kg). Representative images of a) the brain and b) the liver were imaged in IVIS after 24 hours of nanoparticle administered via the tail vein. c) quantification of signal from the liver with and without mannitol of M30 D90.

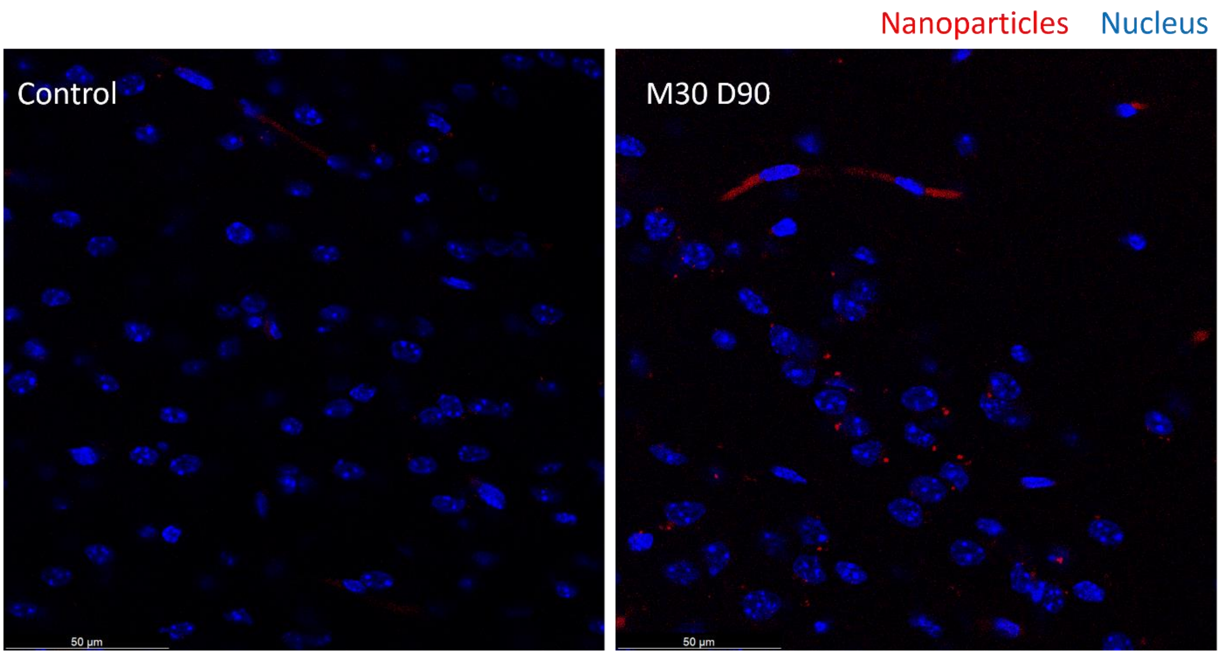


**Figure S8.** In vivo, M30 D90 uptake after 6 hours in caveolae induction model of mannitol. Nanoparticle (red) and Nuclei (blue) (Scale bar:50µm)

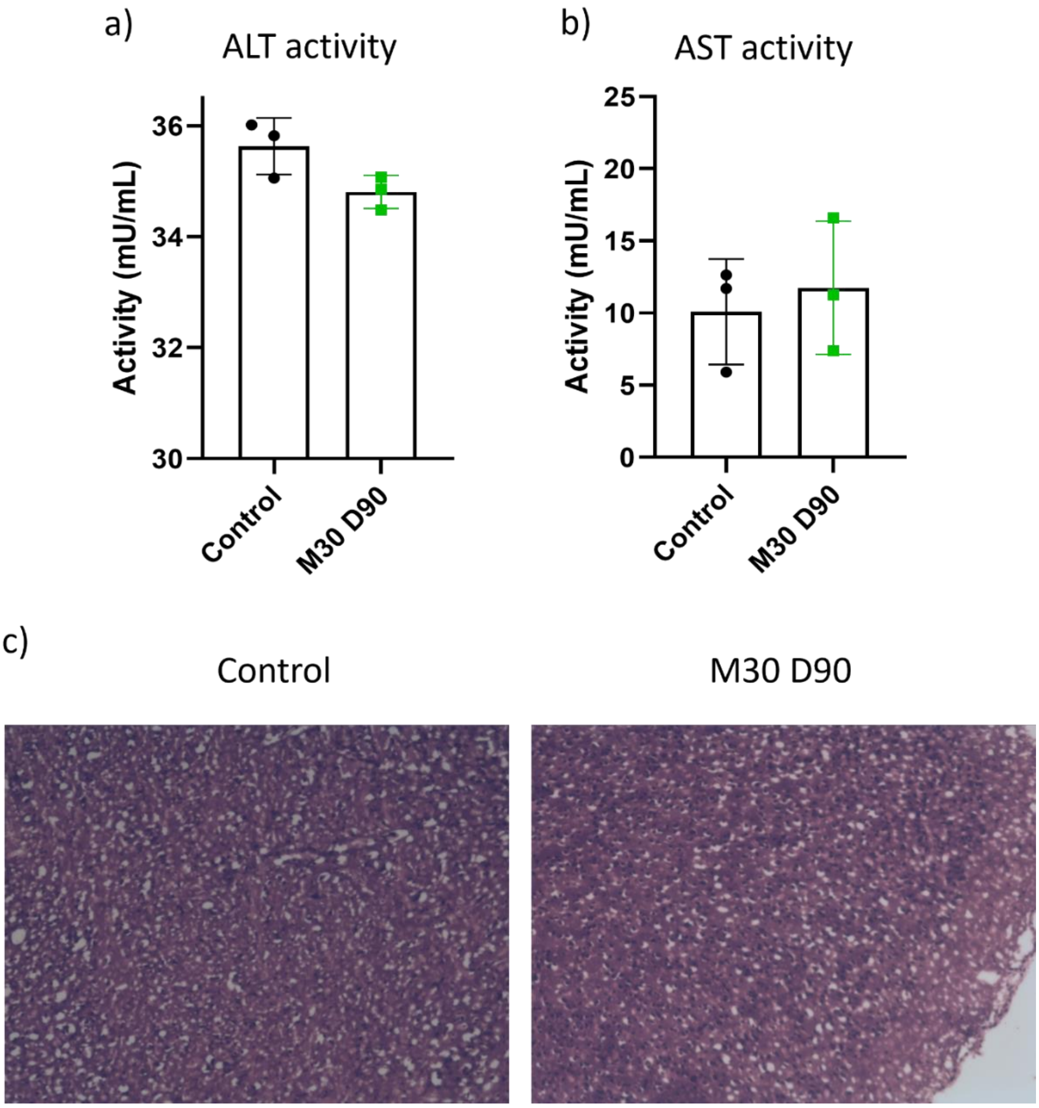


**Figure S9.** In vivo safety profile of M30 D90. Liver enzyme assays were performed with serum collected after 24 hours of M30 D90 injection. a) ALS and b) AST levels of M30 show no elevation to control. c) H&E staining of brain sections.
